# High-Throughput Assessment of Vocal Modulation Following Chemogenetic Inhibition in Songbirds

**DOI:** 10.1101/2025.05.02.651591

**Authors:** Chenyi Song, Mizuki Fujibayashi, Kentaro Abe

**Author notes:** Corresponding author: Contact information Kentaro Abe, Ph. D., Professor, Lab of Brain Development, Graduate School of Life Sciences, Tohoku University. Katahira 2-1-1, Aoba-ku, Sendai, Miyagi 980-8577, Japan. These authors contributed equally.

## Abstract

Chemogenetic tools that enable temporary and repeatable manipulation of neuronal activity have become essential for studying the precise neural underpinning of animal behavior. However, their application has so far been limited to songbirds, which are widely used to study behaviors related to vocal communication. In this study, we applied and evaluated designer receptors exclusively activated by designer drugs (DREADD)-mediated neural suppression in songbird’s brains, focusing on its impacts on vocalizations. We found that neuronal activity in zebra finches (Taeniopygia guttata) can be effectively suppressed both in vitro and in vivo using the inhibitory DREADD by its ligand deschloroclozapine (DCZ). By establishing a high-throughput system for recording and analyzing their vocalizations, we systematically assessed the effects of DREADD-mediated suppression on song behavior. Inhibiting HVC in zebra finches led to a reduction in song production number for approximately 90 min and altered phonological features over a longer time course. Notably, the concentration of DCZ required to induce robust effects on song behaviors was higher than doses typically used in mice. Furthermore, suppression of HVC or Area X in Bengalese finches (Lonchura striata domestica) produced a distinct effect on song features, suggesting species-specific responses to chemogenetic intervention. These findings refine the use of chemogenetic tools in songbirds and provide new insights into the neural mechanisms underlying vocal communication.

## Introduction

Tools for manipulating neuron activity are highly versatile and play a central role in neuroscience studies aimed at uncovering the mechanisms of brain function by linking specific neuronal manipulations to resultant behaviors. Particularly valuable are technologies that enable reversive and repeatable control of defined neuronal populations, thereby allowing precise assessment of their contribution to the behaviors of interest. Moreover, minimally invasive approaches are especially desirable in experiments targeting naturalistic, spontaneous behaviors in freely moving animals. Chemogenetic tools are developed to fulfill these needs, aiming to manipulate neural activity in vivo with the least invasive procedures. Among such tools, the muscarinic acetyl choline receptors that was modified to selectively respond to artificial chemical reagent, namely Designer Receptors Exclusively Activated by Designer Drugs (DREADDs) have been widely employed in the field of behavioral neuroscience.(Armbruster et al., 2007) Since their development, DREADDs have been utilized in a variety of experimental animal species, providing valuable insights into brain function across a wide range of neural processes including behavior, learning and memory, emotion, sensory processing, whole body physiological regulation.(Roth, 2016; Wiegert et al., 2017) The feature of DREADD to allow manipulation on the activity of specific population of neurons and glia within the brain, repetitively and least invasively, is ideal for songbird studies aiming at revealing the neural mechanisms behind communication which necessitates unstressed, natural behavior of subjects.(Fujibayashi and Abe, 2024) However, compared to the widespread usage in other species, the application of DREADD technology in songbirds has been limited to few studies so far.(Heston et al., 2018)

To establish an experimental framework for DREADD-mediated neuronal manipulation in songbirds, this study reassessed the procedures for its application. We focused on hM4Di, which suppresses endogenous activity upon activation, as its effects are more physiologically interpretable and better suited for manipulating brain function (Armbruster et al., 2007). In contrast, other DREADD subtypes, such as hM3Dq and rM3Ds activation enhance neuronal activity but may induce non-physiological excitation patterns, complicating the interpretation of behavioral outcomes.(Alexander et al., 2009) This is particularly important in the context of this study, which aims to evaluate the efficacy of DREADD manipulation in modulating song-related behaviors. We found that the DREADD agonist deschloroclozapine (DCZ),(Nagai et al., 2020) effectively suppressed spontaneous neuronal firing in cultured zebra finch neurons transfected with the inhibitory DREADD, hM4Di, in a manner similar to that observed in mouse neurons. Furthermore, we demonstrated that hM4Di also suppresses neuronal function in vivo in the zebra finch brain, as evidenced by the attenuation of auditory-induced neuronal activation. For their song’s features, we found a reduction in song bout number, and modulation of the phonological features of vocalizations soon after DCZ administration. However, we observed that this effect required approximately tenfold higher DCZ concentrations in vivo compared to those used in rodents. By applying the same procedure in Bengalese finches, we found a similar but distinct effect on song, suggesting a species-specific effect on songs. These findings establish a foundation for employing DREADD technology in songbirds to elucidate the precise neural mechanism underlying interindividual communication.

## Results

### DREADD mediated suppression of neurons in songbird brain

To evaluate the efficacy of DREADD manipulation in the songbird neurons, we prepared dissociated primary neuron cultures from the telencephalon of zebra finch 1 day post hatch. In these cultures, we expressed the inhibitory DREADD receptor, hM4Di, together with the genetically encoded calcium indicator GCaMP6s (Figures 1A and 1B). After 10– 15 days in culture, we applied bicuculline to enhance synaptic transmission among them, which induced periodic fluctuations in the GCaMP6s fluorescent signal driven by spontaneous firing (Figure 1C). These fluctuations were suppressed by tetrodotoxin (TTX, 1 μM) treatment, confirming that they represent excitatory synaptic transmissions among the cultured neurons. The application of DCZ (4.0 μg/mL and 0.4 μg/mL) significantly reduced the fluctuation in GCaMP6s signals. Notably, there was no significant difference in the effect of DCZ treatments between zebra finch and mouse neurons, even at the lowest concentration (Figure 1D; two-way ANOVA, mouse versus zebra finch, *F*(1, 83) = 0.21, *P* = 0.65). DCZ treatment alone caused no effects to neurons without expression of hM4Di (Figure S1). These findings demonstrate that hM4Di effectively suppresses neuronal activity in zebra finch neurons, consistent with its functionality in rodent neurons.

**Figure 1.**
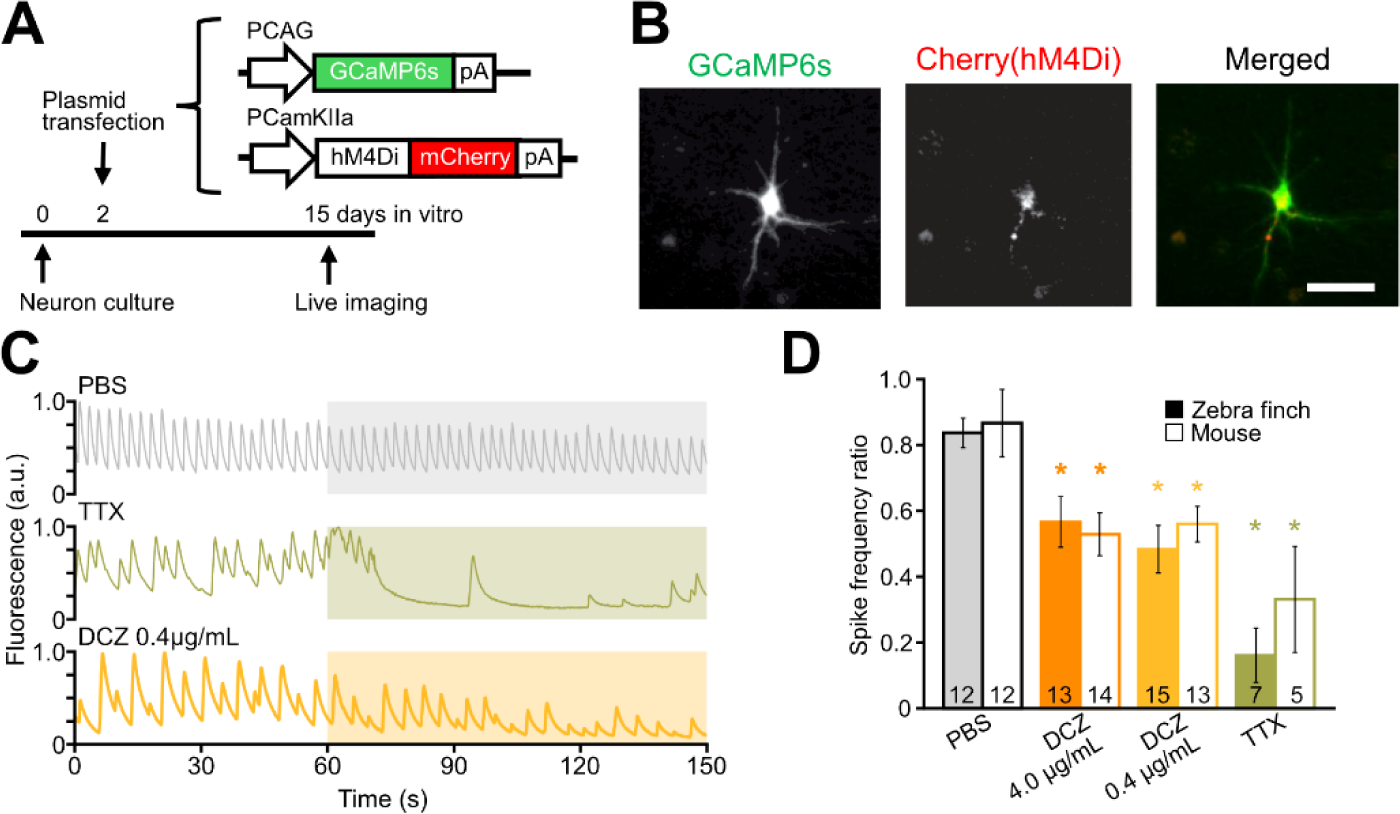
DREADD mediated suppression of neuron activation in cultured songbird neurons *in vitro*. (**A**) Schematic diagram of experimental procedures. Neuron culture from mouse embryo and zebra finch chick was prepared and transfected. (**B**) Fluorescent image of a cultured zebra finch neuron at 15 div. (**C**) Time lapse plot of fluorescent signals from single cells expressing GCaMP6s before and after PBS, TTX, and DCZ application. Drugs are applied to the medium at 60 s (colored highlights). (**D**) Ca^2+^ spike number after drug application (120–180 s) normalized against the value before drug application (0–60 s). * *P* < 0.05 against PBS in each cohort, Dunnett’s test. The number of cells in each cohort are indicated in the bars. Scale bare, 50 µm.

Next, we investigated the impact of DREADD activation on neural activity in vivo. The nidopallium caudomedial (NCM), an auditory region of the songbird brain, is known to exhibit robust neural responses to a playback of conspecific birdsong.(Pinaud and Terleph, 2008) Using an adeno-associated vector (AAV, serotype 2/9 or 2/1), inhibitory DREADD, hM4Di was expressed to neurons within the NCM of zebra finch (Figure 2A). To assess the effectiveness of hM4Di in suppressing neuronal activity, we evaluated auditory responses to conspecific songs and the bird’s own song (BOS) by quantifying Egr-1/ZENK protein expression using immunostaining at 90 min following song presentation (Figure 2B).(Mello et al., 1992) Throughout the experiment, birds were continuously observed by an experimenter nearby to ensure that they refrained from spontaneous vocalization. As anticipated, auditory stimulation elicited a robust induction of Egr-1 expression within NCM (Figure 2C). Intramuscular injections of PBS or DCZ (either 0.1 or 1.0 mg/kg) 60 min before song stimuli, did not result in significant difference in the overall density of Egr-1 positive neurons following song presentation (Figure 2D). However, we observed that DCZ treatment selectively attenuated Egr-1 expression within hM4Di-expressing neurons, indicating effective chemogenetic inhibition (Figure 2E). This suppression exhibited a dose-dependent effect, with significant reduction in Egr-1 expression detected in birds administrated with 1.0 mg/kg of DCZ, but not with 0.1 mg/kg dose (Figure 2E). These findings demonstrate that hM4Di mediated DREADD activation effectively suppresses neuronal activity within the songbird brain in vivo.

**Figure 2.**
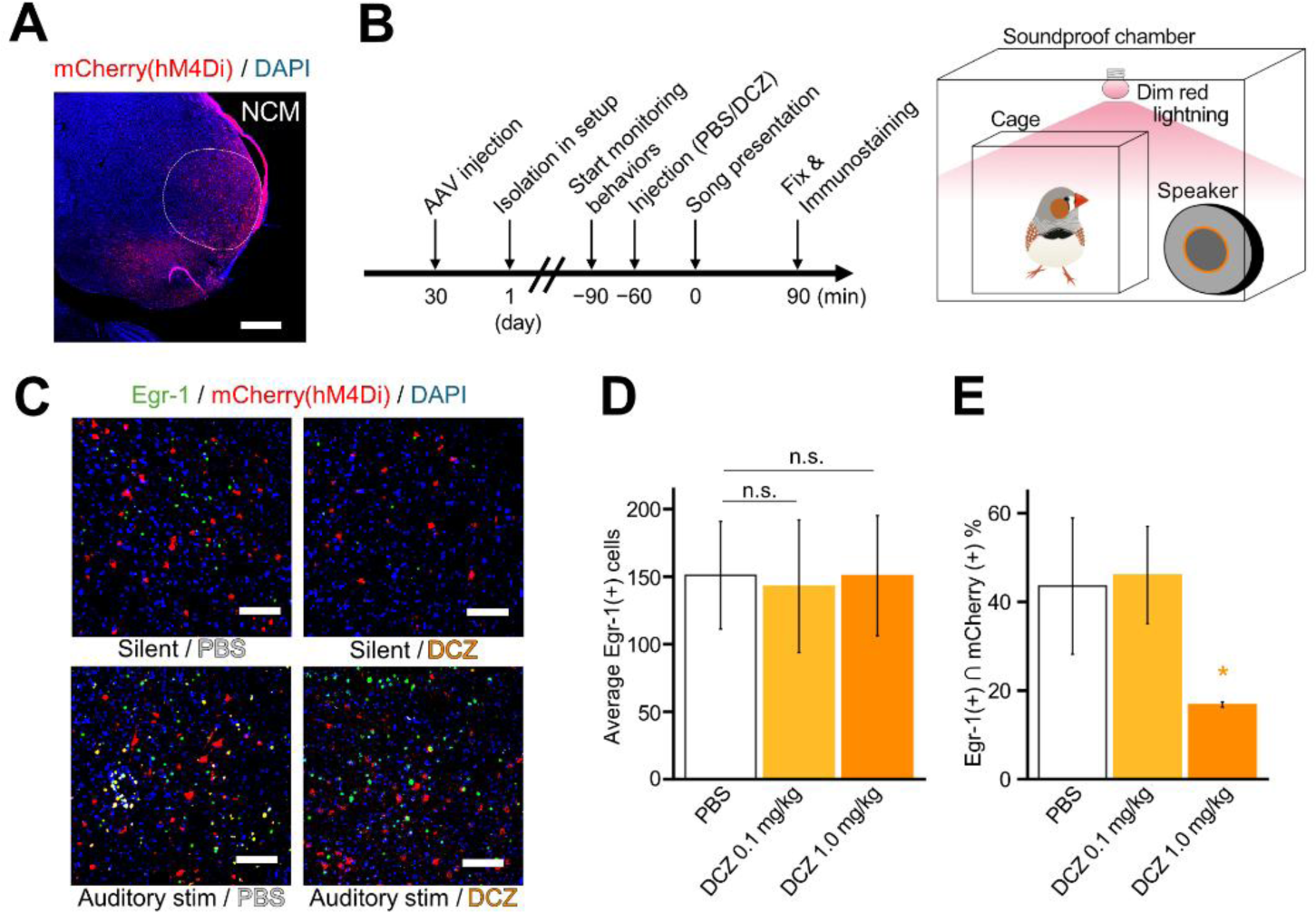
DREADD mediated suppression of auditory responses in songbird brain *in vivo*. (**A**) Immunostained section showing AAV injection site in adult zebra finch brain. NCM, nidopallium caudomedial. Scale bar, 500 µm. (**B**) Schematic diagram of experimental procedures and auditory stimulation setup. (**C**) Immunostained sections. Left, PBS injected bird kept in silence. Right, DCZ injected bird with song presentation. Scale bars, 100 µm. (**D, E**) The number of Egr-1(+) cells (**D**) and Ratio of Egr-1(+) cells within hM4Di (+) cells (**E**) in each group. *n* = 8 sections per group. n. s., *P* > 0.05, * *P* < 0.05 against PBS. *P* values are from the Wilcoxon rank sum test and Bonferroni correction.

### Establishment of the pipeline for systematic analysis of song features

To evaluate the effects of DREADD manipulation on vocal features with particular focus on the time course of their effects, we developed a pipeline for comprehensive song analysis over extended time periods (Figure 3). The bird subjects, injected with an AAV vector to express DREADDs in the target brain nucleus were housed in soundproof chambers and alternately given daily injections of PBS and DCZ over 3–11 cycles (Figures 3A and 3B). Songs were recorded using SAP2011, a standard song detection program in songbird research field.(Tchernichovski et al., 2000) The recorded audio was processed with SAIBS, a machine learning-based system for automatic syllable annotation.(Kawaji et al., 2024) In this study, we performed further customization of SAIBS to extract vocal onset and offset information alongside syllable labels, enabling high-throughput extraction of phonological features of each syllable. Using SAT, a MATLAB interface for SAP2011, this information was exported for downstream analysis (Figure 3C). This integration allowed efficient large-scale analysis of vocal features across a large volume of syllables spanning the entire experimental period. Syllables for phonological analysis were selected from density-based song segments (see STAR Methods). The data was pre-processed using an outlier detection algorithm based on principal component analysis (PCA), after which the phonological features for individual syllables were quantitatively analyzed (Figure 3C). This pipeline allows a systematic and highly efficient framework for evaluating the temporal dynamics of DCZ-induced effects. It facilitates detailed assessment of vocal changes over time, capturing both the immediate consequences of DREADD manipulation and potential long-term post-manipulation plasticity.

**Figure 3.**
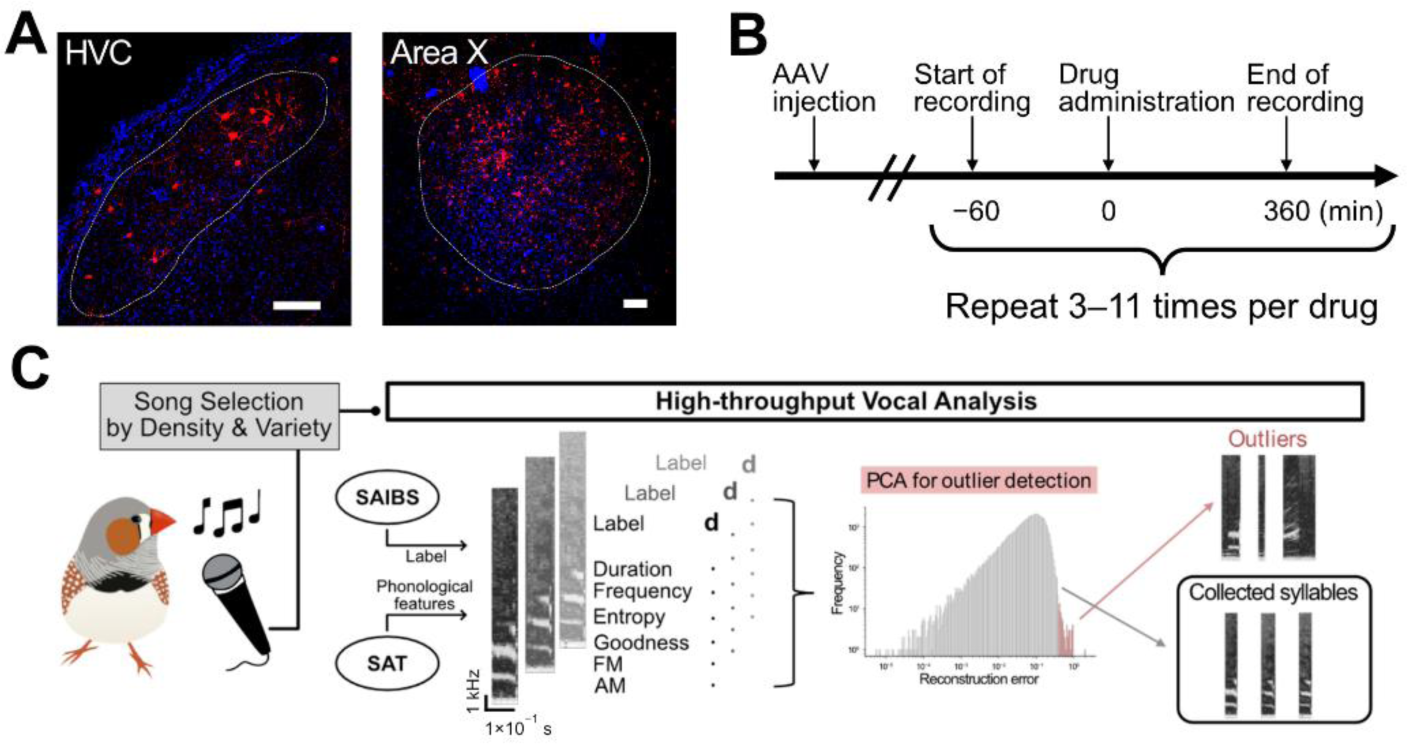
Experimental pipeline for systematic analysis of song features. (**A**) Immunostained sections showing AAV injection site in adult zebra finch brain. Red, hM4Di; blue, DAPI. Scale bars, 100 µm. (**B**) Schematic diagram of experimental procedures. Birds were treated with 150 µL of DCZ or PBS through bilateral pectoral injection around 9:30 a.m. every 1 or 2 days. Their songs were recorded with a microphone and processed using SAP2011 from 60 min before and 360 min after the drug administration. (**C**) Schematic diagram for high-throughput vocal analysis. Syllables from finch songs were annotated by SAIBS and calculated phonological features by SAT. The features produced by SAT were utilized in PCA-based outlier detection to eliminate mislabeled syllables.

### Effect of DREADD mediated suppression on the song features of zebra finches

To investigate the effect of DREADD manipulation on song vocalization, we first focus on the effects on song after suppression of HVC (letter-based name), a brain nucleus playing a central role in song articulation.(Long and Fee, 2008) Previous ablation studies have demonstrated robust and significant deformation and reduction in song bouts, with the severity of the effect depending on the extent of the affected neuronal population.(Chen et al., 2014; Alalawi et al., 2019) To achieve DREADD-mediated suppression of HVC, male birds were bilaterally injected with AAV vectors to express hM4Di in the HVC (Figure 3A). Birds expressing either hM4Di-mCherry or a control construct (mCherry) were isolated in soundproof chambers, and their vocalizations were continuously recorded and analyzed for entire day, throughout the experiment periods (Figure 3B). DCZ or PBS was administered via intramuscular injection at approximately 9:30 AM on recording days. Each subject received alternating injections with PBS and DCZ at approximately 1–2-day intervals, and song features were analyzed within the same individual to allow for within-subject comparison (Figure 4A). In birds expressing hM4Di in HVC, DCZ injection (1.0 mg/kg) led to a transient reduction in song production, with a marked decrease observed during the 0–90 min period following administration (Figure 4B). This reduction was not observed in control birds injected with DCZ, thereby ruling out the possibility that the effect was due to DCZ injection alone. Effect integral over the 0–120 min period, defined as the area between the song production curve and the normalized baseline value, revealed that HVC suppression significantly reduced song production compared to the control birds. At the individual level, the DCZ-induced reduction in song production was consistently observed across eight cycles of DCZ/PBS injections, indicating that the effect is both reversible and reproducible (Figure 4C). The phonological features of the songs were further analyzed using our pipeline for audio feature analysis (Figure 3C). Immediately after the injection of DCZ in HVC suppressed birds, we observed a reduction in duration of motifs, a chunk of stereotypic sequence of syllables (Figures 4D and 4E). As reported previously,(Glaze and Troyer, 2006) natural baseline variations in motif length were observed even in control birds injected with PBS (Figure S2A), but the changes were more pronounced in HVC-suppressed birds after DCZ injection. Because we analyzed motifs with identical syllable sequences, the observed effect was not attributable to changes in the syntactic organization within the motifs, but rather to phonological features of the syllables and the gaps between them. This change in motif length appeared to result from a decrease in syllable length within the motif (Figure S2B).

**Figure 4.**
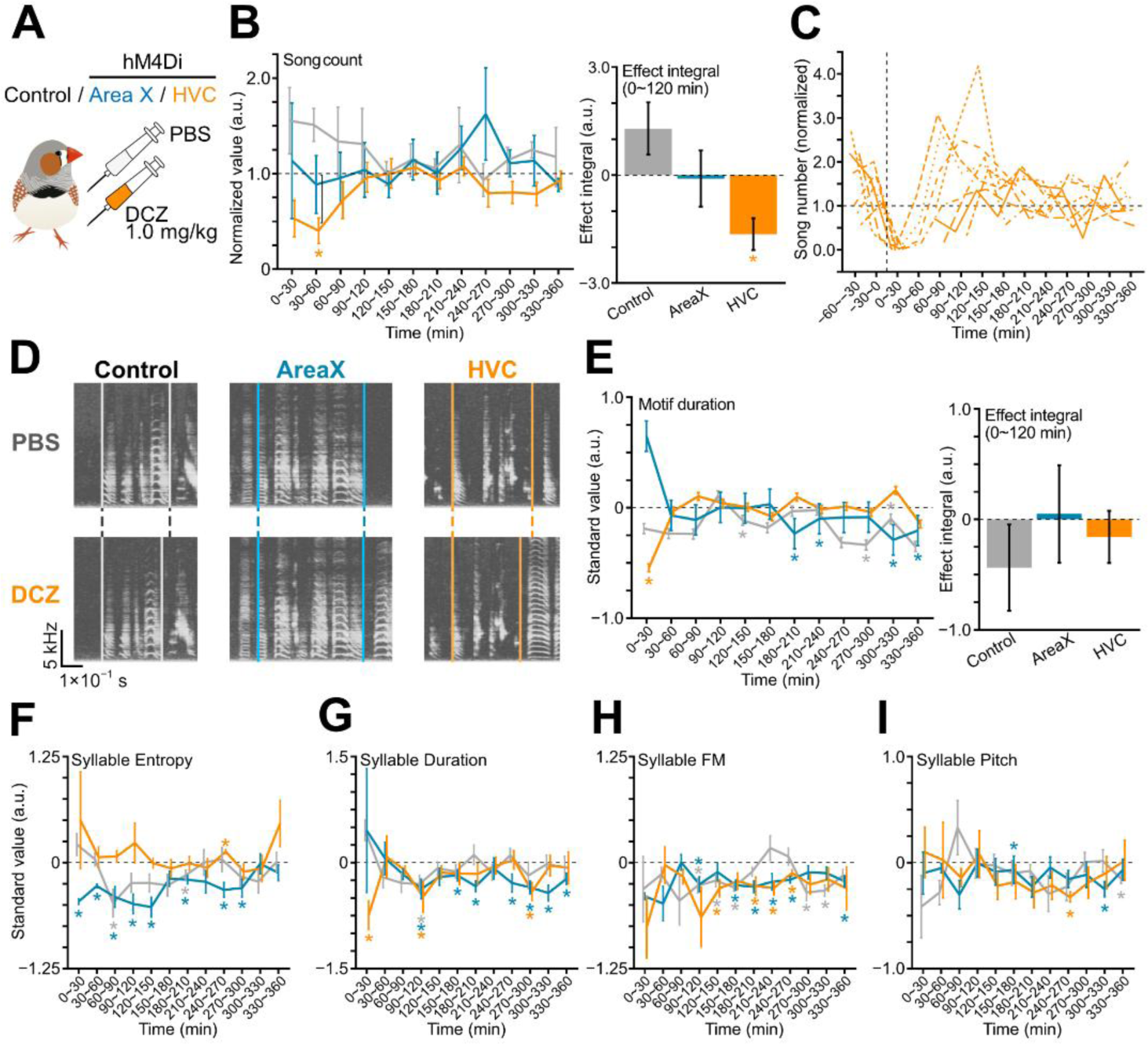
DREADD mediated suppression of HVC and Area X in zebra finches. (**A**) Schematics of experiment. Each day, control birds (black; no hM4Di expression), and birds expressing hM4Di in HVC (orange) or Area X (blue) were injected with PBS or DCZ of 1.0 mg/kg. (**B**) Number of songs produced per 30-min bin, normalized to PBS controls for each corresponding bin. Bar graph on the right displays integrated effects over the 0–120 min period. *n* = 5 (control), 9 (HVC), 7 (Area X) birds. (**C**) Trace showing the number of song bouts across eight DCZ administration cycles in a single subject expressing hM4Di in HVC. (**D**) Representative sonograms from birds injected with PBS / DCZ. For comparison, motifs with the minimum (DCZ) and maximum (PBS) duration are shown to illustrate that changes in motif duration occur without alteration in sequence. Vertical lines indicate the onset and offset of a motif. (**E**) Standardized motif duration. Bar graph on the right displays the integrated effects over the 0–120 min period. *n* = 6 (control), 16 (HVC), 10 (Area X) motifs recorded from 5 (control), 9 (HVC), 7 (Area X) birds. (**F**–**I**) Standardized phonological features of a harmonic syllable within motifs. Wiener entropy (**F**), Syllable duration (**G**), extent of Frequency modulation (**H**), and mean goodness of pitch (**I**). *n* = 8 (control), 11 (HVC), and 12 (Area X) syllables recorded from 4 (control), 7 (HVC), 5 (Area X) birds. Across the panels, values from DCZ-injected days were normalized (**A**, **C**) or standardized (**E**–**I**) to the corresponding PBS-injected values within each 30-min bin. Line and bar plots represent mean ± SEM across subjects. * *P* <0.05. *P*-values displayed in bar graph were calculated using Wilcoxon rank-sum test with Bonferroni correction for multiple comparisons, and those in line graphs using one-sample *t*-test.

Area X is part of the anterior forebrain pathway and is located upstream of LMAN, which is known to play a role in song learning by creating phonological variation in songs.(Fee and Goldberg, 2011) Although Area X is not necessary in adult birds for the motor control in producing songs,(Sohrabji et al., 1990) it is known to provide context dependent modulation of the phonological features of syllables.(Kojima et al., 2018) Contrary to HVC suppression, we observed no significant effect on song production number by DREADD-mediated suppression of Area X (Figures 3A and 4B). In Area X-manipulated birds, we observed a decrease in motif duration at later timepoint after DCZ injection, a phenotype different from HVC-manipulated birds (Figure 4E). We further analyzed the effect on the phonological features, focusing on the harmonic syllable within the motif, which is present in the songs of most individuals.(Price, 1979) As expected from previous studies,(Kojima et al., 2018) suppression of Area X resulted in a persistent reduction in the Wiener entropy of harmonic syllables during the 0–300 min period (Figure 4F). In contrast, this phenotype was not observed in HVC suppressed birds. For both HVC suppression and Area X suppression, we observed significant changes in the length of this syllable (Figure 4G), which generally paralleled the changes in total motif durations, showing a robust decrease immediately after DCZ injection (0–30 min) and a slight but sustained decrease at later time points (180–360 min). We also observed effect on extent of frequency modulation (FM) (Figure 4H), and goodness of pitch at later time points (Figure 4I).

Taken together, the DCZ-induced effect in song features was reversible and reproducible, demonstrating the specificity and consistency of the DREADD-manipulation in songbird brain.

### Dose-dependent effect of DREADD on song vocalization

Next, we evaluated the dose of DCZ required to induce the observed phenotype in zebra finches. For this analysis, we focused on the robust reduction in song number observed in the birds expressing hM4Di in HVC (Figure 4B). We administrated varying doses of DCZ (0.01, 0.1, and 1.0 mg/kg) each day for about 6 rounds, and their effects on song production were assessed compared to vehicle-treated days (Figure 5A). We observed a clear dose-dependent response: no discernible effect at 0.01 mg/kg, a mild reduction at 0.1 mg/kg, and robust suppression at 1.0 mg/kg (Figure 5B). In this experiment, a decrease in song number that reached significant difference was detected during the 30– 60 min periods following DCZ administration at the 1.0 mg/kg dose (Figure 5B).

**Figure 5.**
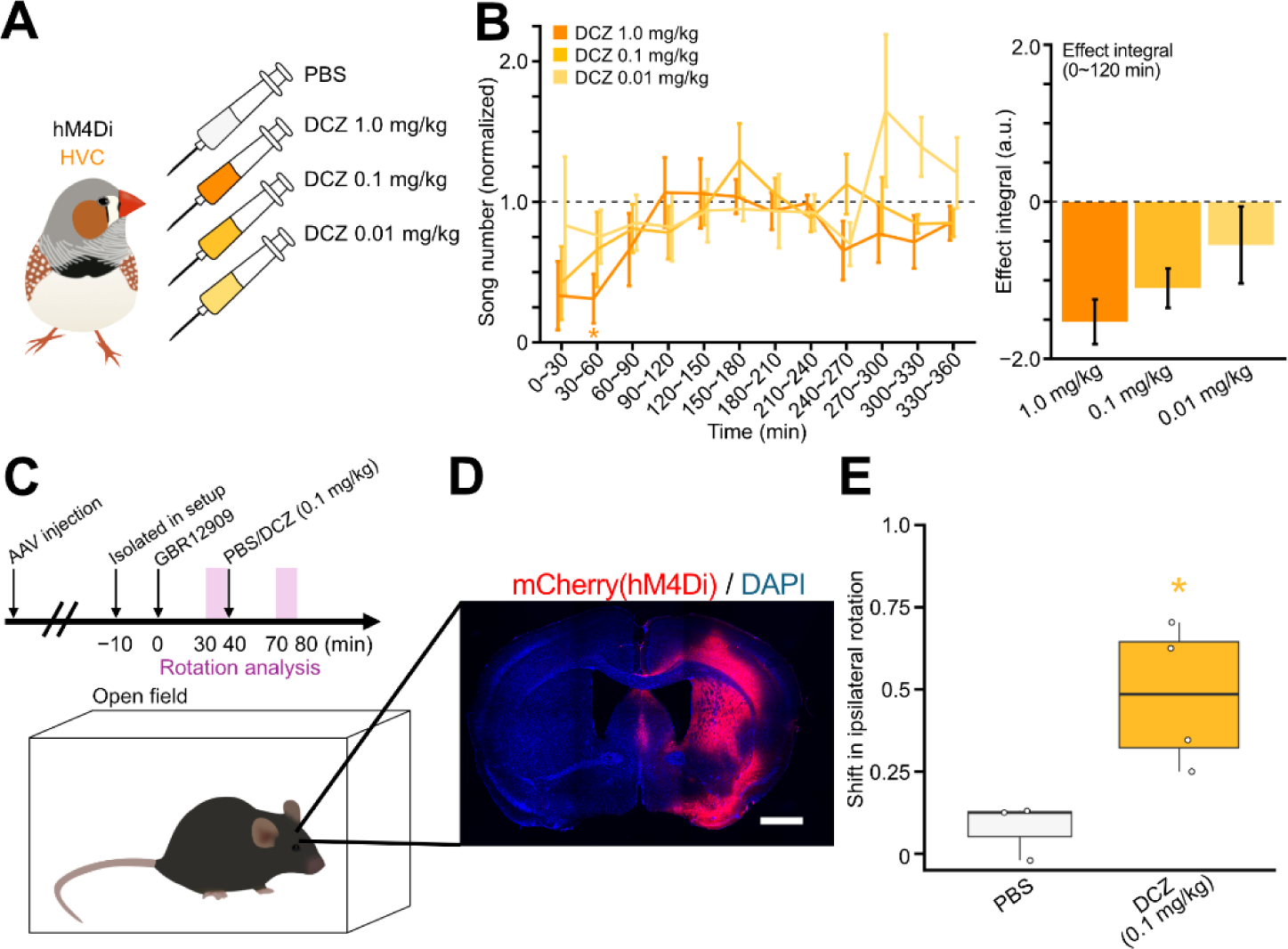
Dose analysis of DREADD effect on songs. **(A)** Schematic diagram of experimental procedures. Zebra finches were treated with three types of concentration DCZ. 1.0mg/kg, 0.1mg/kg, and 0.01mg/kg. 150 µL DCZ solvent was intramuscularly administrated. **(B)** Number of songs produced per 30-min bin, normalized to PBS controls for each corresponding bin. Bar graph on the right displays integrated effects over the 0–120 min period. *n* = 5 birds. * *P* < 0.05. *P*-values were calculated using one sample *t*-test. **(C)** Schematic diagram of experimental procedures and behavioral analysis setup. **(D)** Immunostained brain section of a mouse with unilateral hM4Di injection. Scale bare, 1.0 mm. **(E)** Change in the number of ipsilateral turns toward the hM4Di injected hemisphere following DCZ administration. *n* = 3 (PBS) and 4 (DCZ) treatments were applied to a single mouse. * *P* < 0.05. *P*-values were calculated using Welch’s *t*-test.

We noticed that the effective concentration of DCZ required to elicit robust song-related effects was approximately ten-fold higher than the dose typically used in monkeys and rodents (∼0.1 mg/kg).(Nagai et al., 2020; Nentwig et al., 2022) To confirm the functionality of our hM4Di virus, we transduced the same virus into mouse striatum and analyzed the behavioral effects of DCZ injection (Figures 5C and 5D). Following hemispheric virus injection, we facilitated spontaneous behavior by administering the dopamine reuptake inhibitor GBR12909 (13.3 mg/kg).(Lane et al., 2005) Subsequently, we injected 0.1 mg/kg of DCZ and analyzed behavior around 30–40 min after DCZ administration. We found that DCZ injection (0.1 mg/kg, a suboptimal dose for zebra finches) robustly induced ipsilateral rotation behavior in mice relative to the virus-infected hemisphere as observed in previous studies,(Yu et al., 2022) verifying that our hM4Di viruses function as expected with 0.1 mg/kg DCZ dose in mice (Figure 5E). Taken together, these results suggest that DCZ exhibits a dose-dependent effect in songbirds *in vivo*, requiring a higher concentration to produce robust song-related changes compared to its effect in rodents.

### Effect of DREADD mediated suppression on the song features of Bengalese finches

To further generalize our findings, we applied DREADD-mediated suppression of the HVC and Area X in the Bengalese finch, another songbird species widely studied as a model for vocal communication.(Okanoya, 2004; Sakata et al., 2008; Veit et al., 2021; Kawaji et al., 2024) Compared to the songs of zebra finches, songs of Bengalese finches are relatively longer and more variable in the syntactical organization of syllables.(Okanoya, 2004) We observed no significant difference in the number of cells expressing hM4Di between zebra finches and Bengalese finches (Figures S3A and S3B). In contrast to zebra finches, hM4Di-mediated suppression of HVC with a 1.0 mg/kg dose of DCZ in Bengalese finches did not exhibit significant effect on song production number when compared to the control-treated birds (Figures 6A and 6B). For the phonological features of songs, we observed an immediate reduction (0–30 min) followed by a later extension (180–270 min) of motif duration in both HVC-suppressed and Area X-suppressed individuals (Figures 6C and 6D). Similar to our observations in zebra finches, this effect was not attributable to changes in the syntactic organization within motifs, as our analysis was conducted on motifs with identical syllable sequences. Focusing on harmonic syllables within the motifs, suppression of Area X reduced the syllable’s Wiener entropy immediately following injection (0–30 min) (Figure 6E), a phenotype not observed by HVC suppression. For both HVC suppression and Area X suppression, change in syllable duration (Figure 6F), frequency modulation (FM) (Figure 6G) was observed as with zebra finches although the detailed effect differed between two species. Notably, a steady increase in goodness of pitch was observed around 0–210 min period particularly in Area X suppressed birds (Figure 6H), contrasting to what was observed in zebra finches (Figure 4I). Comparing different features, we observed that phenotypic patterns differed between zebra finches and Bengalese finches (Figure S3C). Together, these findings confirm the effectiveness of DREADD manipulation in another songbird species and suggest species-specific differences in the phenotypic effect on songs.

**Figure 6.**
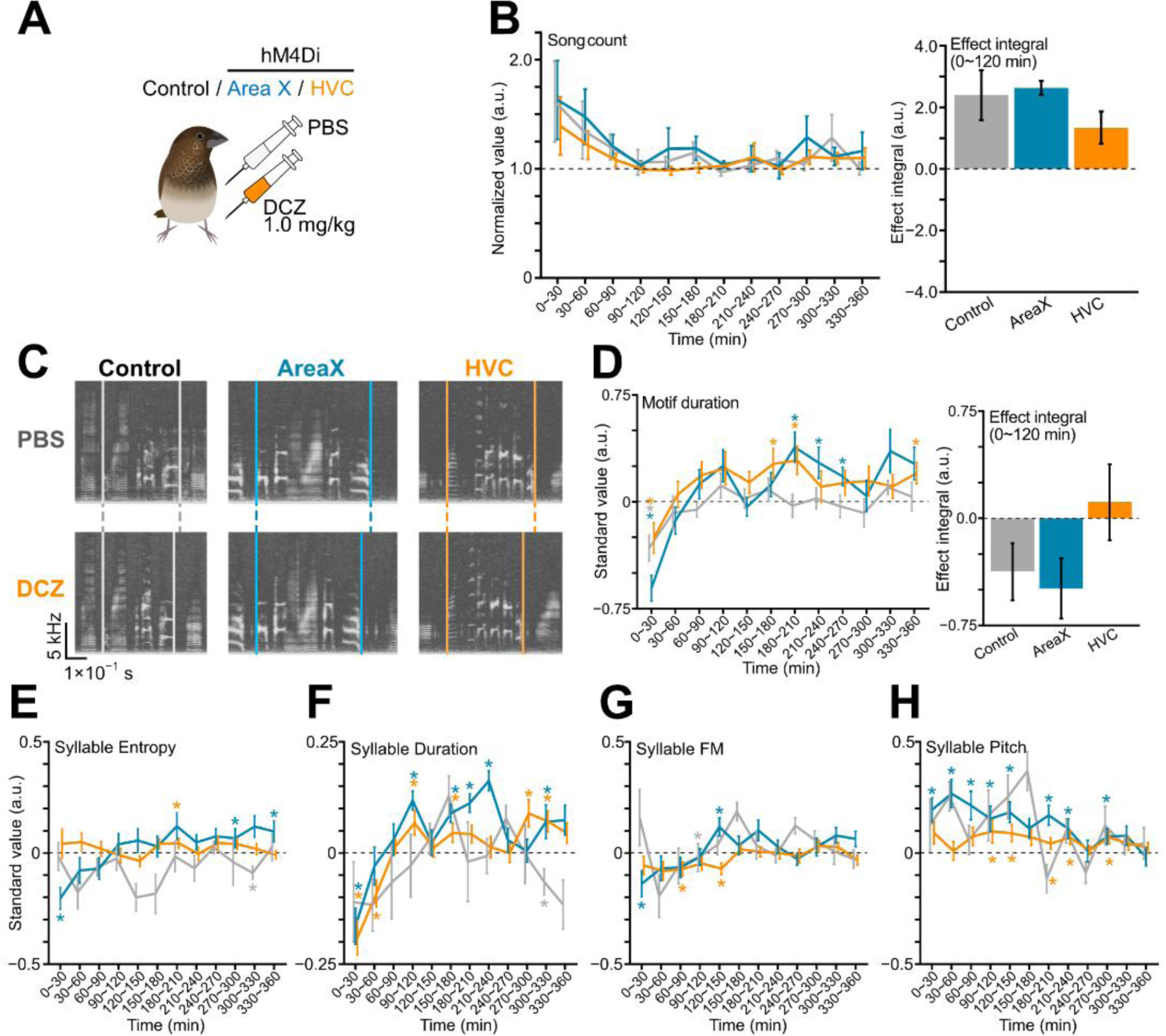
DREADD mediated suppression of HVC and Area X in Bengalese finches. (**A**) Schematics of experiment. Each day, control birds (black; no hM4Di expression), and birds expressing hM4Di in HVC (orange) or Area X (blue) were injected with PBS or DCZ of 1.0 mg/kg. (**B**) Number of songs produced per 30-min bin, normalized to PBS controls for each corresponding bin. Bar graph on the right displays integrated effects over the 0–120 min period. *n* = 4 (control), 5 (HVC), and 3 (Area X) birds. (**C**) Representative sonograms from birds injected with PBS / DCZ. For comparison, motifs with the minimum (DCZ) and maximum (PBS) duration are shown. Vertical lines indicate the onset and offset of a motif. (**D**) Standardized duration of motifs. Bar graph on the right displays the integrated effects over the 0–120 min period. *n* = 8 (control), 12 (HVC), and 7 (Area X) motifs. (**E**–**H**) Standardized phonological features of harmonic syllables within motif. Wiener entropy (**E**), Syllable duration (**F**), Frequency modulation (**G**), and mean goodness of pitch (**H**). *n* = 11 (control), 19 (HVC), 16 (Area X) syllables recorded from 4 (control), 5 (HVC), and 3 (Area X) birds. Across the panels, values from DCZ-injected days were normalized (**B**) or standardized (**D**–**H**) to the corresponding PBS-injected values within each 30-min bin. Line and bar plots represent mean ± SEM across subjects. * *P* < 0.05. *P*-values displayed in bar graphs were calculated using Wilcoxon rank-sum test with Bonferroni correction for multiple comparisons, and those in line graphs using one-sample *t*-test.

## Discussion

DREADD-based manipulation strategies are well-suited for behavioral neuroscience research; however, their application in songbird studies has remained limited. This lack of prevalence suggests potential inefficiencies or failure in practically implementing DREADDs in songbird models. The aim of this study was to identify these limitations and develop improved methodologies for successful application of DREADD techniques in songbirds. In this study, we successfully demonstrated the reversible effects of hM4Di-mediated neuronal suppression in vitro and in vivo. Notably, no differences were observed in DCZ-induced DREADD activation in cultured neurons between mice and zebra finches. However, to elicit phenotypic changes in zebra finch *in vivo*, a DCZ dose of approximately 1.0 mg/kg was required, a dose significantly higher than that commonly used in rodent or primate studies.(Nagai et al., 2020; Nentwig et al., 2022) The lack of observable effects at lower concentrations remains unexplained, but interspecies differences in drug metabolism may contribute to this discrepancy. Recent studies have shown that clozapine-N-oxidate (CNO), the originally developed DREADD ligand,(Bender et al., 1994) exhibits poor brain penetrance and must be metabolized into clozapine to exert its effects.(Gomez et al., 2017) Given that the metabolic profile of CNO varies even among rodent species, such as between rats and mice,(Manvich et al., 2018) it is plausible that the limited success of DREADD application in songbird research may be attributed to species-specific differences in ligand metabolism between avian and mammalian systems. In this study, we used deschloroclozapine (DCZ), a newly developed DREADD ligand shown to exhibit superior brain penetrance compared to CNO in mammalian species.(Nagai et al., 2020) Previous studies have demonstrated that CNO requires approximately 100-fold higher concentrations than DCZ to achieve a comparable effects,(Nagai et al., 2020; Nentwig et al., 2022) suggesting that using CNO in songbirds may require even higher doses than those required for DCZ. We successfully established that a DCZ dose of 1.0 mg/kg is sufficient to induce robust and reversible neuronal suppression in songbirds. While this finding confirms the effectiveness of DREADD-mediated manipulation in songbird models, the overall efficacy of it depends on both adequate receptor expression and ligand pharmacokinetics. Therefore, further investigation into alternative DREADD ligands—such as Compound 21 (Thompson et al., 2018) or JHU37152 and JHU37160 (Bonaventura et al., 2019) —as well as receptors based on different molecular backbones, such as KORDs,(Vardy et al., 2015) may facilitate the development of more efficient and flexible chemogenetic approaches for studying neuronal function in the songbird brain.

Manipulating neural activity in songbird brains is essential for establishing causal relationship between neuronal activity and behavioral outcomes. One of the key advantages of DREADDs is their reversibility and reproductivity. In contrast, traditional approaches used in songbird research have largely relied on irreversible manipulations, such as neuronal ablation via local injection of ibotenic acid,(Sohrabji et al., 1990; Foster and Bottjer, 2001) caspase activation,(Sánchez-Valpuesta et al., 2019) or chromophore targeted ablation.(Scharff et al., 2000) While effective, these non-reversible methods often lack experimental flexibility and reproducibility within individual subjects, necessitating larger sample size to obtain robust results. Reversible approaches have also been explored in songbirds, including the use of tetrodotoxin (TTX) or GABA_A_R agonists,(Cardin and Schmidt, 2004; Andalman and Fee, 2009; Hamaguchi and Mooney, 2012) and optogenetic tools(Xiao et al., 2018; Singh Alvarado et al., 2021; Katic et al., 2022) to transiently silence neuronal activity. However, these methods typically require invasive procedures, such as brain surgeries to implant cannulas or optical fibers for the delivery of pharmacological agents or light. Compared to these methods, another significant advantage of DREADD technology is its minimal invasiveness during experimental manipulation of targeted neurons. Unlike traditional approaches, the DREADD-based manipulation requires only a single viral transduction procedure performed well in advance of the experimental sessions. Subsequent manipulation of neuronal activity can be achieved through rapid and minimally invasive methods, such as intramuscular injection or oral administration of the ligand. These features make DREADDs particularly well-suited for studies of natural, undisturbed behaviors such as free vocal communication in songbirds, where minimizing stress and experimental disruption is essential.

By applying DREADD-mediated suppression in the songbird brain, we observed both similarities and differences in experimental outcomes, compared to those reported using conventional methodologies.(Chen et al., 2014; Alalawi et al., 2019; Isola et al., 2020) One unexpected finding was the effect on motif and syllable duration observed in both zebra finches and Bengalese finches, which had not been robustly observed in previous studies using pharmacological suppression. The discrepancy may, in part, stem from differences in the specific neural target manipulated by each approach, and the extents of off-target effects. For instance, whereas methods such as ablation or TTX application affects both excitatory and inhibitory neurons, hM4Di expression in our study might restricted to specific neuron types because of the use of CaMKIIα promoter which express transgenes more specifically in excitatory neurons.(Dittgen et al., 2004; Scheyltjens et al., 2015) This feature of the DREADD methodology, the ability to identify the manipulated population in a post hoc manner, provides an advantage over conventional approaches, in which the targeted cells are often difficult to visualize. For example, the use of specific promoters enables selective manipulation of defined neural populations, and single-cell sequencing can be used to confirm the identity of the manipulated cells.(Colquitt et al., 2021) Another possible contributing factor is the temporal nature of the manipulation. Conventional methods often involve chronic or irreversible modulation, while DREADD-based approaches allow for acute and reversible modification, which is often reported to produce distinct phenotypes.(Otchy et al., 2015) Particularly, we observed differences in the time-course of song phenotypes between HVC- and Area X-manipulated birds. Specifically, the effect of HVC manipulation occurred relatively rapidly, within 120 min following DCZ injection, whereas the effect of Area X manipulation tended to persist to later timepoints. Since DREADD effects in mammals are generally immediate,(Nagai et al., 2020) the mechanism underlying this delayed effect remains unclear. One possible explanation is homeostatic plasticity, whereby birds adjust their vocalization in response to perturbation.(Konishi, 1965; Tumer and Brainard, 2007; Andalman and Fee, 2009) Because Area X is not considered to be directly required for vocal production but instead contributes to plastic changes in their songs,(Fee and Goldberg, 2011) this may underlie the difference in time course of song phenotype between HVC and Area X manipulation. These possibilities were not directly addressed in this study, and further work will be required to determine the cause of time differences in song phenotype.

We observed differential effects on song phenotypes following DREADD mediated neuronal suppression between zebra finches and Bengalese finches. Although the overall number of hM4Di-expressing cells did not differ significantly between the two species, the resulting effects on song phenotype were distinct. Further investigation is required to determine whether these differences arise from variation in the affected cell populations; nonetheless, they may reflect species-specific roles of HVC and Area X in song vocalization. Notably, the HVC in Bengalese finches demonstrates auditory feedback response during singing,(Okanoya and Yamaguchi, 1997) a feature absent in zebra finches.(Hamaguchi et al., 2014) The absence of an effect on song production frequency in Bengalese finches following DREADD-mediated suppression of HVC— contrasting with the reduction observed in zebra finches—may reflect species specific differences in the functional organization of HVC. In particular, Bengalese finches may perceive discrepancies between intended and actual vocal output, potentially allowing them to compensate for disrupted motor commands. Alternatively, differences in the intrinsic connectivity within the HVC may also contribute to this variation. Neurons within the HVC exhibit highly stereotyped sequential activity during song production, driven by synfire chain networks formed through interconnections among these neurons.(Hahnloser et al., 2002; Long et al., 2010) It has been hypothesized that Bengalese finches possess branching, parallel synfire chains,(Jin, 2009) which may compensate for the impact of suppressing a small subset of neurons. In contrast, zebra finches are thought to rely on more liner synfire chains, making their vocal output more vulnerable to neuronal suppression occur. These functional difference in HVC neurons may be established by substantial differences in their gene expression profiles between the two species.(Kato and Okanoya, 2010) Further studies are needed to reveal the species-specific mechanism of song control and to explore the evolutional origins of these differences.

### Limitations of the study

As a limitation of study, the potential off-target effects of DCZ in avian species remain insufficiently explored. In our experiments, administration of DCZ at a dose of 1.0 mg/kg alone did not produce significant changes in song phenotypes in control birds, in contrast to the effect observed in Area X- or HVC-manipulated birds. However, in certain song features, similar trends were observed across all groups during the early post-administration period (0–90 min periods; Figures 6B and 6D). Further detailed analysis of whole-body physiology after DCZ injection, or carefully designed control experiments, will be necessary, considering the molecular and genetic differences between mammals and birds. The observed 10-fold differences in the dose required for DREADD activation in vivo between mammals and birds suggest the need for more efficient brain-penetrant ligands for use in avian species. Furthermore, it remains unclear whether the difference in the effects of HVC or Area X suppression observed in this study compared to those using non-reversible methods were due to differences in acute or chronic suppression. Further studies utilizing similar chemogenetic tools designed for non-invasive and rapid manipulation of neuronal function will be required to address this question and refine our understanding of the mechanism of song control.

## Acknowledgments

This work was supported by Tohoku University Research Program “Frontier Research in Duo” No.2101, JSPS/MEXT KAKENHI JP25H01037, JP24H01218, JP24H02146, JP23K18252, JP21K19424 awarded to K.A. and JP25KJ0541 awarded to M.F. We thank the members of the Abe Lab at Tohoku University for help and fruitful suggestions.

## Author contributions

C.S., M.F. and K.A. conceived and initiated the project. K.A. performed cell culture experiments and data analysis. C.S. performed *in vivo* experiments and data analysis. M.F. developed the methodology, performed experiments, and data analysis. C.S., M.F. and K.A. wrote the initial manuscript. All authors discussed and commented on the manuscript.

## Material and Methods

### Animals and Cares

The care and experimental manipulation of mice and birds used in this study were reviewed and approved by the institutional animal care and use committee of Tohoku University. All experiments and maintenance were conducted following relevant guidelines and regulations.(Olson et al., 2014) Birds were purchased from Asada Chojyu and kept in our aviary under a 14-hour light / 10-hour dark cycle (daytime: 8:00−22:00); water and food were given *ad libitum*.

### Cell culture and Ca^2+^ Imaging

Primary neuron culture from zebra finches (*Taeniopygia guttata*) were prepared from individuals of post hatch day 1. The forebrain of their brain was collected and processed with enzymatic digestion with papain (Roche, #10108014001; 300 U/mL) for 20 min at 37°C. The tissues are titrated to dissociate into single cells and plated onto 24-well plates pretreated with poly-L-lysin (SIGMA-Aldrich, #P2336). Cells are cultured in DMEM/F12 (Fujifilm-Wako) with 10% fatal bovine serum (FBS, Biowest) for initial plating, and then maintained with an equal mixture of DMEM/F12 with 10% FBS and Neurobasal plus medium (Thermo-Fisher, #21103049) supplemented with B27 plus supplement (20×; Thermo-Fisher, #A3582801), Glutamax-I (100×; Thermo-Fisher, #35050061) and penicillin-streptomycin (100×; Fujifilm-Wako) at 37°C under 5% CO_2_. Primary cortical neuron culture from mouse (*Mus musculus*, Slc:ICR) was prepared as described in previously.(Abe and Abe, 2022; Yamamoto et al., 2025) In brief, the cortex was collected from embryo at embryonic day 15 and processed with 0.05% Trypsin (Fujifilim-Wako, #201-18841). The cultured mouse neurons are maintained with Neurobasal plus medium supplemented with B27 plus supplement, Glutamax-I, and penicillin-streptomycin. Both zebra finch and mouse neurons were transfected with plasmids using HilyMax reagent (Dojindo, #H357) at 2 days *in vitro* (div). Live imaging was performed at 10–15 div at 37°C, using inverted fluorescence microscope (Axiovert A1 MAT, Zeiss) equipped with a Colibri 7 system (Zeiss) and a stage warmer (Tokai hit). Approximately 30 min prior to the imaging, bicuculline (bicuculline methiodide 200 μM; Tocris, #131) was applied to the well. Fluorescent signals were captured at 10 Hz using 20× objective lens. A droplet of DCZ (4.0 or 0.4 μg/mL in 20 μL PBS; Deschloroclozapine dihydrochloride, MedChemExpress, #HY- 42110A) or tetrodotoxin (TTX, 1.0 μM, Nakarai-tesque, #32775-51) were applied during the imaging at specified timings. Spike numbers from GCaMP6s signals of each neuron were extracted by PCA/ICA method(Mukamel et al., 2009) and Detection of raising event algorithms of Mosaic analysis software (Inscopix).

### Surgery and DCZ application

Adult zebra finches, Bengalese finches and mice (*Mus musculus*, C57BL/6J) were anesthetized with medetomidine-midazolam-butorphanol mixture (Medetomidine 30 mg/mL, All Japan pharma; Midazolam 30 mg/mL, Astellas Pharma; Butorphanol tartrate 500 mg/mL Meiji Seika Pharma; NaCl 118 mM; 100 µL per bird and 350 µL per mouse).(Yamamoto and Abe, 2022) In zebra finches, AAVs (1.0 µL per bird) are injected at the coordinate from the Y sinus: anterior, 0 mm; lateral, 2.2 mm; depth, 0.75 mm at beak angle 40° for HVC; anterior, 0.5 mm; lateral, 0.6 mm; depth 1.0 mm and 0 mm; lateral, 2.2 mm; depth, 2.0 mm at beak angle 40° for NCM; and anterior, 5.5 mm; lateral, 1.2 mm; depth, 3.1 mm and 2.9 mm at beak angle 60° for Area X. In Bengalese finches, AAVs (1.0 µL per bird) were injected at the coordinate from the Y sinus: anterior, 5.75 mm; lateral, 1.2 mm; depth, 2.9 mm and 2.7 mm at beak angle 60° for Area X; and anterior, 0 mm; lateral, 2.0 mm; depth 0.75 mm at beak angle 40° for HVC. For a mouse, AAVs (1.0 µL per mouse) were injected unilaterally into the right striatum at six locations with the coordinate from bregma: anterior, 0 mm; lateral, 2.0 mm; depth, 3.0 mm, 2.6 mm and 2.3 mm; and anterior, 1.0 mm; lateral, 1.8 mm; depth, 3.0 mm, 2.6 mm and 2.3 mm. DCZ/PBS application was performed more than 30 days after the injection. 10 mg/mL stock solution of deschloroclozapine-hydrocloride (DCZ-2HCl) was diluted at desired concentration with 150 μL PBS per bird and injected into the pectoral muscle around 9:30 AM every 1–2 days.

### Viruses and plasmid constructs

Plasmid used for primary neuron transfection are following: pAAV-CAG-GCaMP6s (RRID: Addgene_100844, a gift from Douglas Kim & GENIE Project),(Chen et al., 2013) and pAAV-CaMKIIa-hM4Di-mCherry (RRID: Addgene_50477, a gift from Bryan Roth). AAVs used for *in vivo* transduction are following: AAV2/9-CaMKIIa-hM4D(Gi)-mCherry (titer ≥ 1.0×10¹³ vg/mL, Addgene viral prep #50477-AAV9; 1.0 µL per bird) for hemispheric injection into NCM in auditory response experiments; 4:1 mixture of AAV2/9-CaMKIIa-hM4D(Gi)-mCherry, and AAV2/9-Syn-EGFP (3.54×10¹³ DNase resistant particle/mL); or AAV2/1-CaMKIIa-hM4D(Gi)-2a-EGFP (1.1×10¹³ DNase resistant particle/mL) and AAV2/9-Syn1-mCherry (1.8×10^14^ DNase resistant particle/mL); or AAV2/9-Syn1-mCherry (1.31×10¹³ DNase resistant particle/mL) and AAV2/9-Syn1-EGFP (3.54×10¹³ DNase resistant particle/mL) for bilateral injection at Area X and HVC for song phenotype analysis; 4:1 mixture of AAV2/9-CaMKIIa-hM4D(Gi)-mCherry, and AAV2/9-Syn1-EGFP for mouse striatum. AAVs are obtained from Addgene, or produced in house as reported previously.(Abe and Abe, 2022)

### Auditory stimulation and immunostaining

Auditory stimulation was conducted on birds >30 days after AAV injection. Male birds were isolated in a soundproof chamber and exposed to a randomized playback of a mixture of their own song and conspecific songs (5–9 s for a bout) randomly every minute for 30 min. Subjects were sacrificed 90 min after the auditory stimulation. To prevent vocalization of songs and calls before and after the auditory stimulation, the chamber was illuminated with dim red light, and an experimenter continuously monitored the birds near the chamber to ensure they neither sang nor fell asleep. The birds were rapidly euthanized after 90 min and perfused with PBS followed by 4% paraformaldehyde (PFA, Nakarai-tesque) in PBS. The cryo-sections were immunostained with Egr-1 (1:800; SantaCruz, #sc-189) and RFP (1:1000; MBL, #M165-3MS) with Alexa-488 or Alexa-555-conjugated secondary antibodies (1:450; Thermo-Fisher, #A121422 and #A11008), followed by 4′,6-diamidino-2-phenylindole (DAPI, 0.33 ng/mL, TCI) staining. Images were taken by BZ-810 or BZ-9000 (Keyence) using 4× lens or 10× lens and the whole section merged images were used for the inspection of the infected area, or by FV4000 (Olympus) using 4×, 10×, and 20× lens, and the Z-projection images were created using cellSens FV (Olympus). The expression cell was detected by visual inspection of the stained images using OpenCV-Python (UK Intel Corporation). We used a single section (8 sections per group) containing the highest number of EGR-1 expressing cells. After binarization and watershed processing, hM4Di- or Egr-1-expressing cells were identified based on the circularity and area of signal clusters in each channel. Co-expression was defined when the distance between the centers of signal clusters across channels was shorter than the sum of their estimated radius, in which case the cells were classified as hM4Di and Egr-1 co-expressing cells.

### Song recording and analysis pipeline

Songs were recorded using SAP2011(Tchernichovski et al., 2000) and processed with SAIBS(Kawaji et al., 2024) to classify and annotate syllables, as well as to extract vocal onset and offset time along with syllable labels. Song segments were extracted according to changes in the proportion of time that syllables occurred per unit time and the diversity of syllable types within high-density regions. From these extracted song segments, syllables were selected for subsequent phonological analysis. We utilized SAT, a MATLAB interface of SAP2011 to extract vocal features between the vocal onset and offset of each syllable. To exclude miss-annotated syllables, the data was then pre-processed through an outlier detection algorithm using principal component analysis (PCA). For each syllable type, we conducted PCA using the major vocal features of each syllable: duration, average frequency, average frequency modulation, average amplitude modulation, average goodness, and average entropy. After transforming the data, retaining the principal components that explain more than 90% of the variance, the inverse transformation was performed. Then, we calculated the mean squared error (MSE) between the reconstructed data and the original data. Using these MSE scores, anomaly detection was performed using the Smirnov-Grubbs test at a significant level of 0.05. The resultant phonological features for each syllable were analyzed. This outlier detection was conducted using Python.

### Analysis of songs

Songs were counted every 30 min for 60 min before and 360 min after the injection of PBS/DCZ. For each individual, song counts on DCZ-injected days were normalized to the mean count on PBS-injected days. Motifs were defined as stereotyped sequences of syllables that consistently appear in zebra finch songs, and as invariant sequences appearing with a frequency of at least once per song in Bengalese finch songs. Motif duration was calculated as the duration between the onset of the first syllable and the offset time of the last syllable. All phonological features extracted by using the pipeline mentioned in Figure 3 were standardized by subtracting the mean PBS value and dividing by the standard deviation of PBS values for each 30-min time bin. For zebra finch data, outliers were excluded from each time bin using the interquartile range method following standardization. Effect integral was calculated as the area between the line graph and the baseline value of 1 for song counts, and 0 for motif durations. As controls, we prepared subjects expressing control constructs exclusively in either HVC or Area X, but the subjects were pooled together for analysis.

### Effect of DREADD inhibition on mouse behavior

Mouse behavioral analysis of DREADD induced hemispheric suppression of the striatum was conducted in an open field box (40 × 40 × 40 cm) constructed from gray plastic plates. The subject with hemispheric injection of hM4Di virus 35 days in advance, was habituated to the box for 10 min before the test. GBR12909 (13.3 mg/kg in 200 μL of 20% DMSO/PBS, i.p., TCI #G0541) was administered first, followed by either DCZ (0.1 mg/kg in 100 μL PBS, i.p.) or vehicle (100 μL PBS, i.p.) at 40-min intervals. All drugs were administered into the right inguinal region. DCZ and PBS treatments was administered as a consecutive-day pairs, with a 1–2-day interval before repeating the cycle, resulting in three repetitions of each treatment. Behavior was video recorded at a resolution of 480 × 640 pixels and 30 fps using a USB-camera positioned above the open-field box. From the recorded video, the number of right and left turns was manually counted from 30–40 min after drug administration. A rotation was defined as a movement in which the subject turned more than 180° within a circle with a diameter approximately equal to the body length, without pausing for 1–2 s.

### Quantification and statistical analysis

The alpha score of 0.05 was used to reject the null hypothesis. Two-way ANOVA with post hoc Dunnett’s test, Wilcoxon’s rank sum test, and Bonferroni correction were used for multiple comparisons. An independent sample *t*-test was used for the comparison of two data sets. One sample *t*-test was used for the comparison of one dataset and the baseline value. Statistical analyses were conducted using R.

## Supplementary information

**Figure S1.**
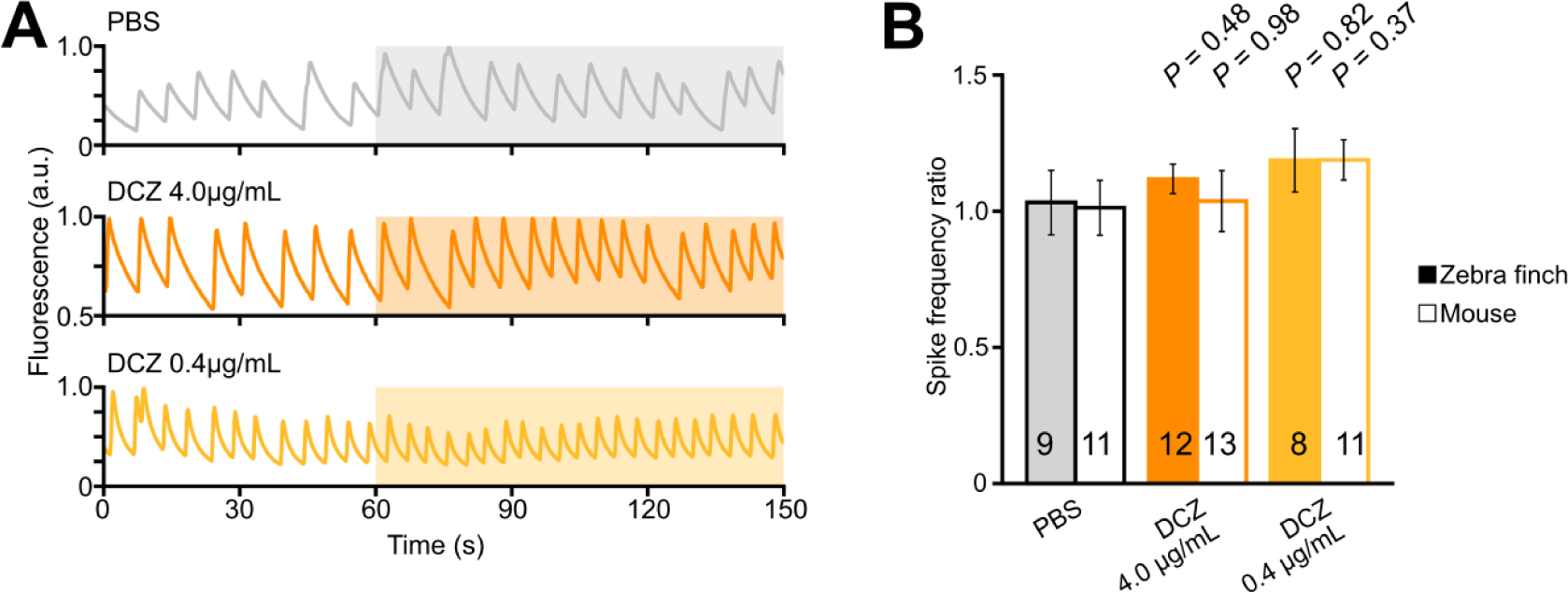
Effect of DCZ on cultured neurons without DREADD receptors. **(A)**Time lapse plot of fluorescent signals from single cells expressing GCaMP6s before and after PBS and DCZ application. Drugs are applied to the medium at 60 s (colored highlights). (**B**) Ca^2+^ spike number after drug application (120–180 s) normalized against the value before drug application (0–60 s). *P* values shown were calculated by Dunnett’s test compared against PBS in each cohort. The number of cells in each cohort are indicated in the bars.

**Figure S2.**
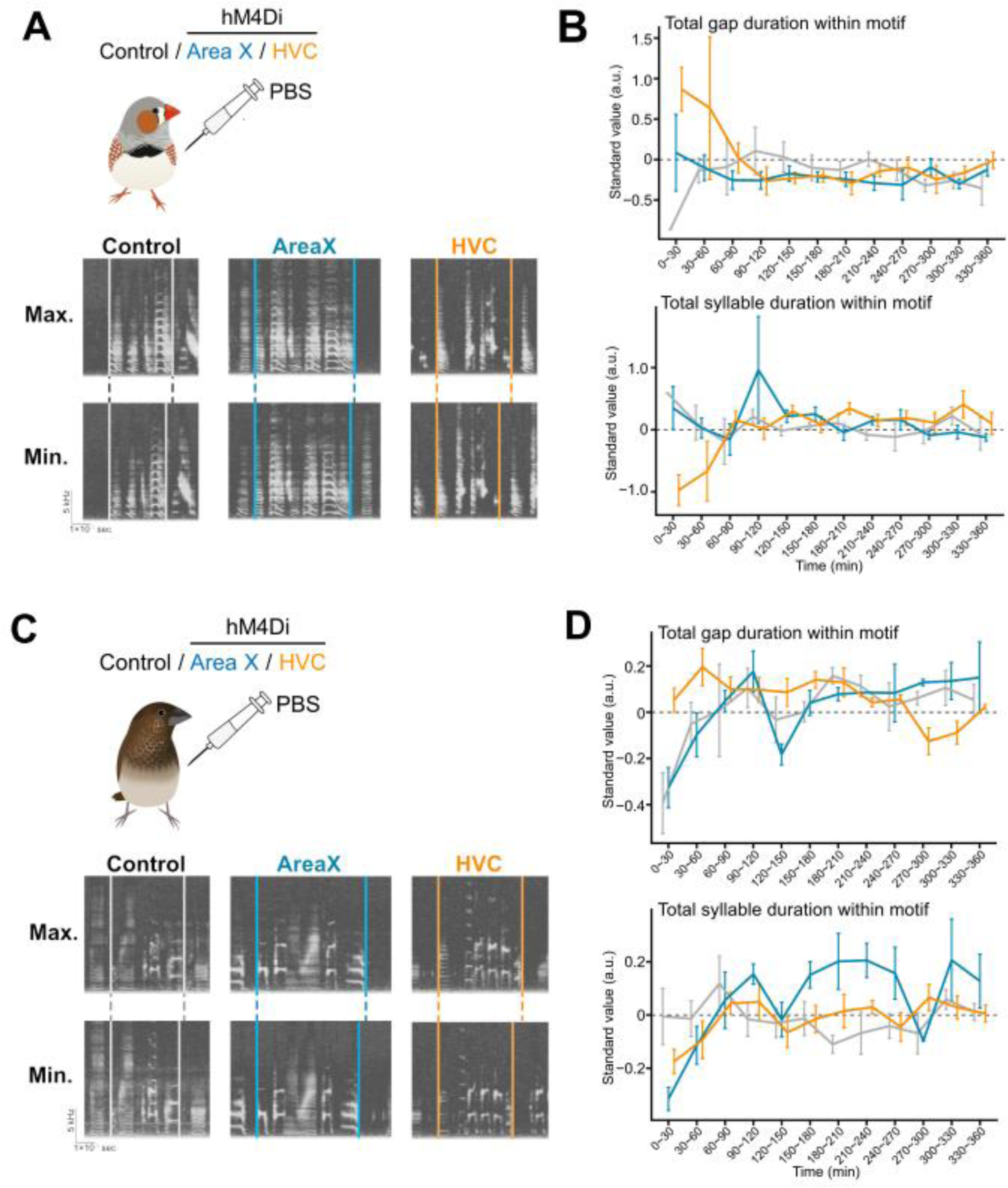
Additional data on song phenotypes. (**A**, **C**) Natural variation of motif length in zebra finch (**A**) and Bengalese finch (**C**) expressing mCherry (control), hM4Di in Area X (blue) or HVC (orange). Sonograms showing the maximum (Max.) and minimum (Min.) motif duration from birds injected with PBS are shown. Vertical lines indicate the onset and offset of a motif. (**B**) Total gap (top) and syllable (below) duration within motif. Zebra finches, n = 7 (control), 16 (HVC), 9 (Area X) motifs. (**D**) Same as (**B**) in Bengalese finches, n = 8 (control), 12 (HVC), 7 (Area X) motifs.

**Figure S3.**
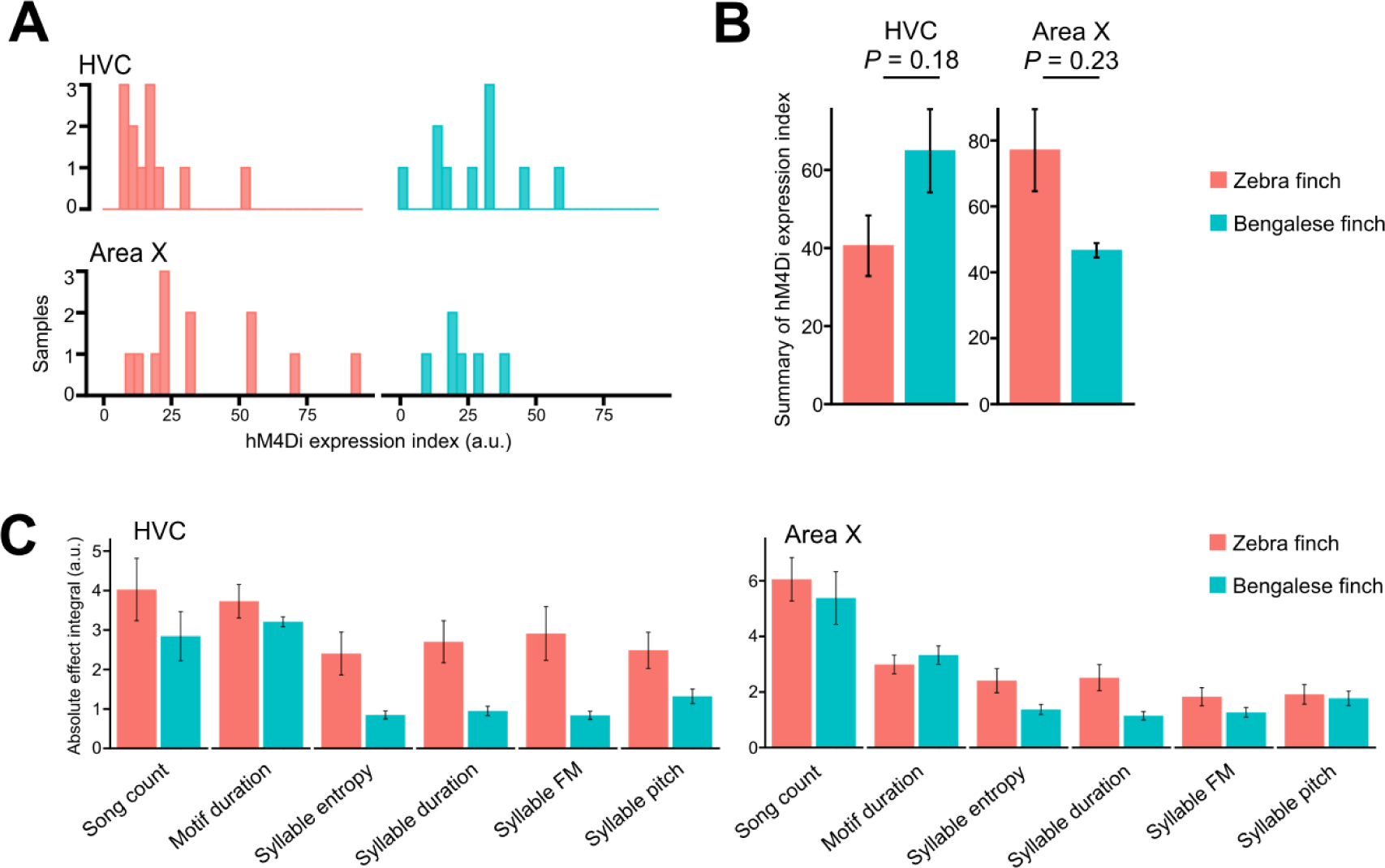
Evaluation of DREADD expression and song phenotype between two species. (**A**, **B**) Quantification of hM4Di expression in the birds used for song phenotype analyses. The number of hM4Di-expressing cells in a single hemispheric brain section was counted for birds injected in HVC and Area X (**A**), and the bilateral summary of those counts for each bird (**B**). Number of birds analyzed: HVC-manipulated birds—zebra finches, 6; Bengalese finches, 5. Area X-manipulated birds—zebra finches, 5; Bengalese finches, 3. Mean ± sem. *P*-values displayed were calculated using Weltch’s t-test. (**C**) Summarized graph showing the effects on six song features—song count, motif duration, syllable entropy, syllable duration, syllable FM, and syllable pitch—in zebra finches and Bengalese finches. Data presented in Figures 4 and 6 are summarized and shown. The absolute area under curve during the 0–360 min period after DCZ injection is shown. Mean ± sem.

